# Brain dynamics predict oral contraceptive treatment duration

**DOI:** 10.64898/2026.08.17.745238

**Authors:** Anira Escrichs, Yonatan Sanz-Perl, Gustavo Deco, Belinda Pletzer

## Abstract

Oral contraceptives are used by millions of women worldwide, yet their cumulative effects on the female brain remain poorly understood. We analyzed resting-state fMRI from 192 women (never, current, and past users) using brain-dynamics metrics that quantify information transfer across spatiotemporal scales. Duration of use was associated with a progressive, age-independent modulation of brain dynamics: prolonged use was related to restricted local synchronization but enhanced information flow across scales. Machine learning predicted individual duration of use from brain dynamics, and node-level classifiers distinguished users from never-users based on a spatially distributed cortical pattern spanning multiple functional networks, most clearly detectable in longterm users. In past users, these associations were inverted in sign, scaling with prior duration rather than returning to the never-user baseline, suggesting active reorganization following cessation. Together, these results indicate that cumulative oral contraceptive exposure acts as a neuromodulator of information processing.

## Introduction

Combined oral contraceptives (COCs) contain synthetic estrogen and progestin analogues that exert their biological effects through estrogen and progesterone receptor signaling and suppress the production of endogenous ovarian hormones (Pletzer et al., 2023). Oral contraceptive use has been related to gray matter volumes in areas crucial for cognitive and emotional processing, including the prefrontal cortex, anterior cingulate cortex (ACC), fusiform gyri, hippocampus, and basal ganglia (Pletzer et al., 2019, 2015). Long-term use has been associated with structural differences in the hippocampus and amygdala (Brønnick et al., 2020). At the functional level, resting-state connectivity has been related to oral contraceptive use, particularly within the default-mode network (DMN), salience, and control networks (Hidalgo-Lopez et al., 2023a; Casto et al., 2022; Petersen et al., 2014). These findings have used complementary methodological approaches: binary group comparisons between users and non-users, duration-dependent changes in long-term users (Noachtar et al., 2022; Pletzer et al., 2023), as well as placebo-controlled trials (Hidalgo-Lopez et al., 2023a; Haase et al., 2025). Connectivity metrics range from seed-based analyses of specific regions such as the amygdala and ACC (Hidalgo-Lopez et al., 2023b), to effective connectivity analyses of large-scale brain networks (Hidalgo-Lopez et al., 2023a), and multivariate brain connectivity analyses (Haase et al., 2025). However, these metrics fail to capture the complex spatiotemporal dynamics of brain communication.

Advances in theoretical neuroscience have established turbulent-like dynamics as a mathematical framework for quantifying how information propagates across spatiotemporal scales, analogous to the energy cascade in fluid turbulence (Deco and Kringelbach, 2020; Deco et al., 2025). Brain turbulence metrics measure how brain activity is organized across short-range versus long-distance connections. This multiscale approach has proven effective for distinguishing macroscale brain states (Escrichs et al., 2022) and measuring reconfigurations in response to pharmacological treatments (Escrichs et al., 2025). Furthermore, dense-sampling longitudinal fMRI data have shown that brain dynamics fluctuate across the natural menstrual cycle (Pritschet et al., 2020), and that the initiation of oral contraceptive use blunts these rhythmic transitions, leading to a constrained turbulent dynamical regime (De Filippi et al., 2021). While these N-of-1 findings established the sensitivity of brain dynamics to synthetic steroids, they cannot characterize long-term cumulative effects across a broader population. Extending these metrics to a larger cohort and coupling the results with machine learning offers a promising approach for decoding individual biological signatures directly from resting-state fMRI (Escrichs et al., 2024).

In this study, we investigated the cumulative effects of combined oral contraceptive use on whole-brain dynamics in a cohort of 192 women (never, current, and past users). We applied the turbulent dynamics framework to resting-state fMRI data to test whether COC treatment duration relates to brain information processing in current users. Machine learning was then used both to predict individual exposure duration from brain dynamics and to classify never-users from oral contraceptive users at the nodal level across spatial scales. Finally, we examined whether these dynamical associations persisted or reorganized following contraceptive cessation in past users.

## Results

### Participants

The study included 192 women: 54 current COC users (age: 22.1 ± 2.7 years, duration: 4.7 ± 2.6 years, range 0.08–11.0), 78 past users (age: 25.1±4.3 years; prior duration: 4.1±3.4 years, *N* = 74), and 60 never-users scanned in the follicular phase (age: 22.7 ± 3.4 years). The correlation between age and duration was strong in current users (*r* = 0.676, *p < .*001) and weaker in past users (*r* = 0.320, *p* = 0.005).

### Overview of the framework

We applied the turbulent-dynamics framework to test whether COC duration is associated with changes in brain information processing across spatial scales. The framework quantifies information propagation across spatial scales using the Kuramoto local order parameter as a spatiotemporal indicator of synchronization (Deco and Kringelbach, 2020; Escrichs et al., 2022). The scale inverse-parameter *λ* controls coupling distance: high values (*λ* = 0.30, ≈3 mm) capture local short-range dynamics, while low values (*λ* = 0.01, ≈100 mm) capture global long-range integration. We derived measures across 11 scales (*λ* ∈ {0.30, 0.27, . . ., 0.01}): Turbulence (variability of local synchronization); Information Transfer (spatial propagation of synchronization at a given scale); Information Cascade Flow (cross-scale information propagation over time); and Information Cascade (average cascade flow across all scales). Together, these metrics provide complementary characterizations of how information is transferred across spacetime scales (see Methods for a detailed description).

### Oral contraceptive duration reorganizes brain information processing across scales

We fitted linear regression models (Metric(*λ*) ∼ Duration + Age) across all spatial scales *λ*. Duration yielded consistent, scale-dependent reorganization (Figure 1), while age was not a significant predictor in any model (*p >* 0.05). Brain Turbulence showed a biphasic pattern across spatial scales (Figure 1a): prolonged use was negatively associated with local-scale turbulence (*λ* ≥ 0.24; *β* = −0.533 at *λ* = 0.30, *p* = 0.004) and positively associated with global-scale turbulence (*λ* ≤ 0.09; all *p <* 0.030), indicating reduced local synchronization variability alongside increased long-range integration. The transition occurred near *λ* ≈ 0.15, where the duration coefficient crosses zero, suggesting a spatial equilibrium point in the dynamic regime. Information Transfer was negatively associated with duration across 10 of 11 scales (Figure 1b; *λ* = 0.30–0.03; *p* ranging from *<* 0.001 to 0.019), with the largest effect at the most local scale (*λ* = 0.30: *β* = −0.648, *p <* 0.001), reflecting a duration-dependent suppression of local-to-intermediate information propagation. Information Cascade Flow was positively associated with duration across all scales (Figure 1c; *λ* = 0.27–0.01; all *p <* 0.012; peak *β* = 0.617 at *λ* = 0.15), and the global Information Cascade confirmed this shift (Figure 1d; *β* = 0.595, *p <* 0.001). The opposing directions of Transfer and Cascade Flow indicate a progressive reconfiguration of multiscale communication architecture: local propagation is suppressed while cross-scale hierarchical flow is enhanced with cumulative COC exposure.

**Figure 1:**
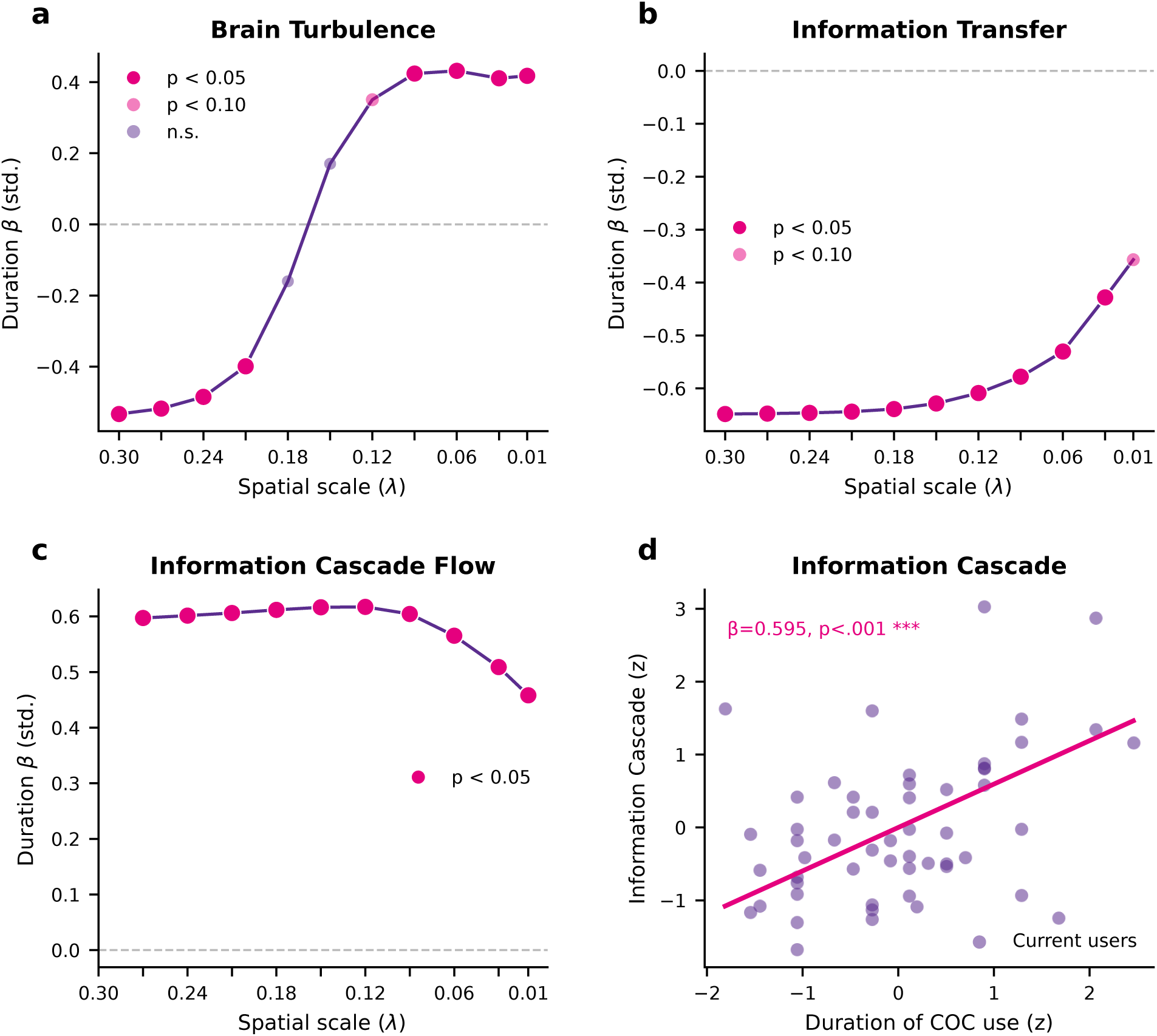
Scale-dependent reorganization of brain dynamics with oral contraceptive duration. All regression models are controlled for chronological age. **(a)** Biphasic effect of duration on Brain Turbulence: decreased local synchronization (*λ* ≥ 0.24) and increased global integration (*λ* ≤ 0.09). **(b)** Spatial Information Transfer decreases with longer oral contraceptive use across 10 of 11 scales. **(c)** Information Cascade Flow increases with duration across scales. **(d)** Global Information Cascade is significantly associated with duration (*β* = 0.595, *p <* 0.001), indicating a time-dependent upregulation of hierarchical integration. Magenta circles indicate significance at *p <* 0.05.

### Brain dynamics predict individual oral contraceptive duration

To evaluate the predictive power of brain dynamics as a marker of contraceptive exposure, we employed a nested cross-validated Lasso regression framework to decode individual duration of use. The scale-level predictive analysis revealed a differential sensitivity of the dynamic metrics to the duration of COC use (Figure 2). Brain Turbulence (Figure 2a) shows predictive capacity predominantly at short-distance spatial scales (*λ* = 0.24 to 0.30, *p* ≤ 0.028) and at global scales (*λ* = 0.01 to 0.09, *p* ≤ 0.024), while middle spatial scales (*λ* = 0.12, 0.15, 0.18) did not show a generalizable predictive gain over the age-only baseline model (*p >* 0.1). Information Transfer (Figure 2b) exhibited the highest robustness among all metrics, yielding significant predictive results across nearly all analyzed scales. This sensitivity suggests that prolonged COC exposure induces a reorganization in the brain’s capacity for spatial communication rather than targeting isolated functional clusters. The maximum predictive gain (Δ*r* ≈ 0.11) was observed at short-range scales (*λ* = 0.30, *p <* 0.001), indicating that the most discriminative changes are anchored in the immediate neighborhood of neural signaling. Information Cascade Flow (Figure 2c) showed a consistent pattern of significance (*p* = 0.018) across all analyzed *λ* scales, reinforcing the evidence of a fundamental reorganization in the hierarchical propagation of information. Finally, the Global Information Cascade (Figure 2d) significantly outperformed the age-only baseline model (Δ*r* ≈ 0.09, *p <* 0.001), establishing global hierarchical integration as a predictive signature of cumulative COC exposure.

**Figure 2:**
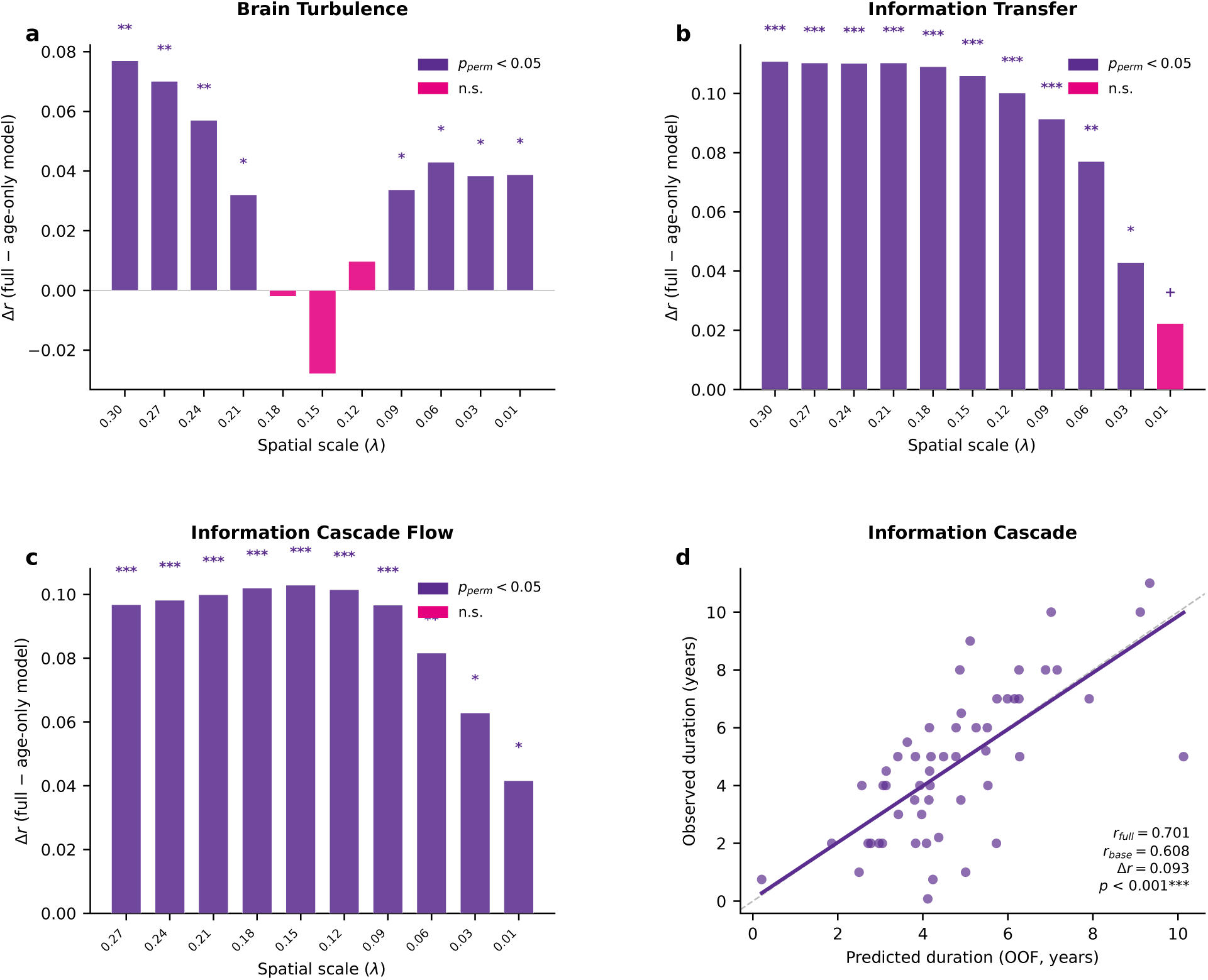
Brain dynamics predict individual oral contraceptive duration across spatial scales. All models use a 10-fold cross-validation framework with Lasso regression. Predictive gain (Δ*r*) is defined as the difference in out-of-fold Pearson correlation between the full model (metric + age) and an age-only baseline, validated via 1000 permutation tests. Significance markers: +*p <* 0.10, ∗*p <* 0.05, ∗ ∗ *p <* 0.01, ∗ ∗ ∗*p <* 0.001. Purple bars indicate *p*_perm_ *<* 0.05; magenta bars indicate non-significant scales. **(a)** Predictive gain of Brain Turbulence, significant predominantly at short-range (*λ* = 0.24–0.30) and global scales (*λ* = 0.01–0.09). **(b)** Predictive gain of Information Transfer, significant across nearly all scales and peaking at local spatial scales (Δ*r* ≈ 0.11, *λ* = 0.30). **(c)** Predictive gain of Information Cascade Flow, significant across all analyzed scales. **(d)** Scatter plot of observed versus out-of-fold predicted duration (years) for the global Information Cascade. The full model (*r*_full_ = 0.701) significantly outperforms the age-only baseline (*r*_base_ = 0.608; Δ*r* ≈ 0.09, *p <* 0.001).

### Oral contraceptive exposure is decodable across spatial scales

To localize this signature anatomically, we trained node-level classifiers (Random Forest, RF; XGBoost, XGB) on 1033 features per participant (1000 nodal turbulence values plus 33 global turbulence-framework descriptors) to separate never-users from COC users, with age residualized within each cross-validation fold and significance assessed by label permutation (1000 permutations; age-residualized ROC AUC). Performance was first mapped across the full scale sweep (*λ* = 0.30–0.01) as a descriptive landscape and then tested at four reference scales spanning the hierarchy (short, *λ* = 0.30; middle, *λ* = 0.18; transitional, *λ* = 0.15; global, *λ* = 0.03; Figure 3).

**Figure 3:**
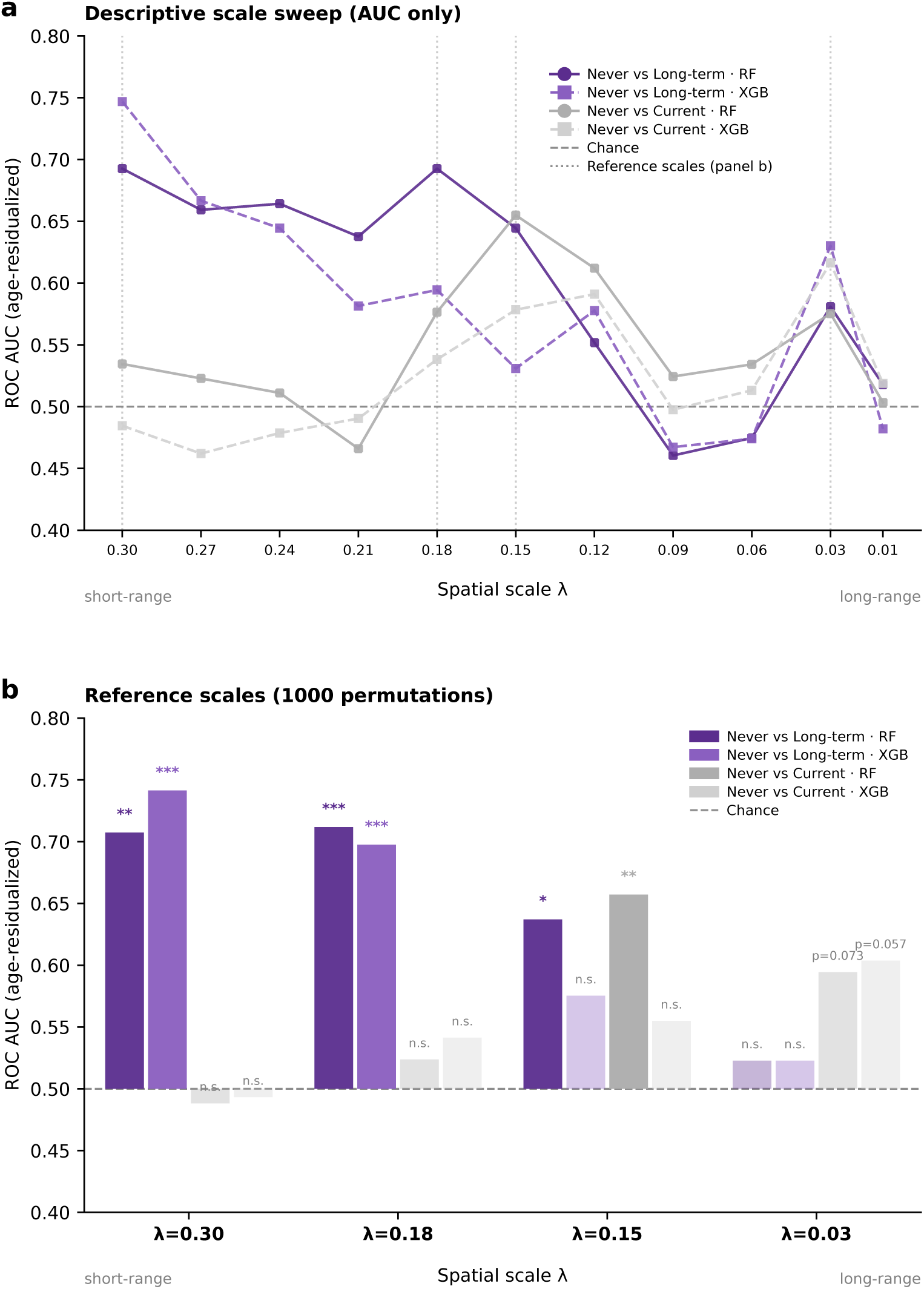
Oral contraceptive exposure is decodable across spatial scales. Node-level classification (1033 features: 1000 nodal turbulence values plus 33 global descriptors) separating never-users from COC users, age-residualized within fold, with label-permutation significance (1000 permutations; age-residualized ROC AUC). **(a)** Descriptive scale sweep across all scales (*λ* = 0.30–0.01; AUC only, no inference), motivating the four reference scales. **(b)** Inferential classification at the four reference scales (*λ* = 0.30, 0.18, 0.15, 0.03) for never vs current and never vs long-term (≥ 4.75 y), Random Forest (RF) and XGBoost (XGB). Long-term users are robustly separable at short and middle scales (AUC up to 0.74); current users emerge only at the transitional scale *λ* = 0.15 (RF AUC = 0.66). Significance markers: ∗*p <* 0.05, ∗ ∗ *p <* 0.01, ∗ ∗ ∗*p <* 0.001; n.s., not significant.

Considering all current users irrespective of duration (*N* = 54), separability was at chance at short and middle scales (*λ* = 0.30: AUC = 0.49, *p* = 0.51; *λ* = 0.18: AUC ≤ 0.54, *p* ≥ 0.30), indicating that COC use *per se* leaves no detectable short- or middle-range signature. Current users became separable precisely at the transitional scale *λ* = 0.15 for RF (AUC = 0.66, *p <* 0.01), while the more strongly regularized XGB remained at chance (*p* = 0.24); at the global scale *λ* = 0.03 both classifiers showed a strong trend (RF AUC = 0.59, *p* = 0.073; XGB AUC = 0.60, *p* = 0.0579). In contrast, long-term users (duration ≥ 4.75 y, the cohort median; *N* = 27) were separable from never-users at short and middle scales (*λ* = 0.30: AUC = 0.71 RF, *p <* 0.01, and 0.74 XGB, *p <* 0.005; *λ* = 0.18: AUC = 0.71 RF and 0.70 XGB, both *p <* 0.005), remained separable for RF at *λ* = 0.15 (AUC = 0.64, *p* = 0.026), and collapsed to chance at the global scale (*λ* = 0.03: AUC = 0.52, *p* ≥ 0.38). The duration signature is thus confined to short- and middle-range dynamics, whereas the current-use trace emerges at the transition between short- and long-range scales.

### Cortical importance maps of the predictive signatures

To interpret which features drove classification, we computed mean per-node feature importance within each of the eight cortical networks (normalized for network size) together with the 33 global features, and the direction of the underlying group difference (mean in COC users minus never-users), at every scale where a comparison was significant, using the best-performing significant classifier at that scale.

For current users (Figure 4), the signature at the transitional scale *λ* = 0.15 (RF) was spatially distributed across networks rather than focal, and the most important single feature was a DMN node in the right medial prefrontal cortex (RH DefaultA PFCm), followed by further DMN and dorsal-attention nodes. At the network level, the DMN, dorsal-attention, temporoparietal, and somatomotor systems ranked highest, with reduced nodal turbulence among users relative to never-users across most of the top features. The global-scale profile (*λ* = 0.03; XGB, *p* = 0.0579) shifted to a control/precuneus and DMN emphasis, reported as a strong trend.

**Figure 4:**
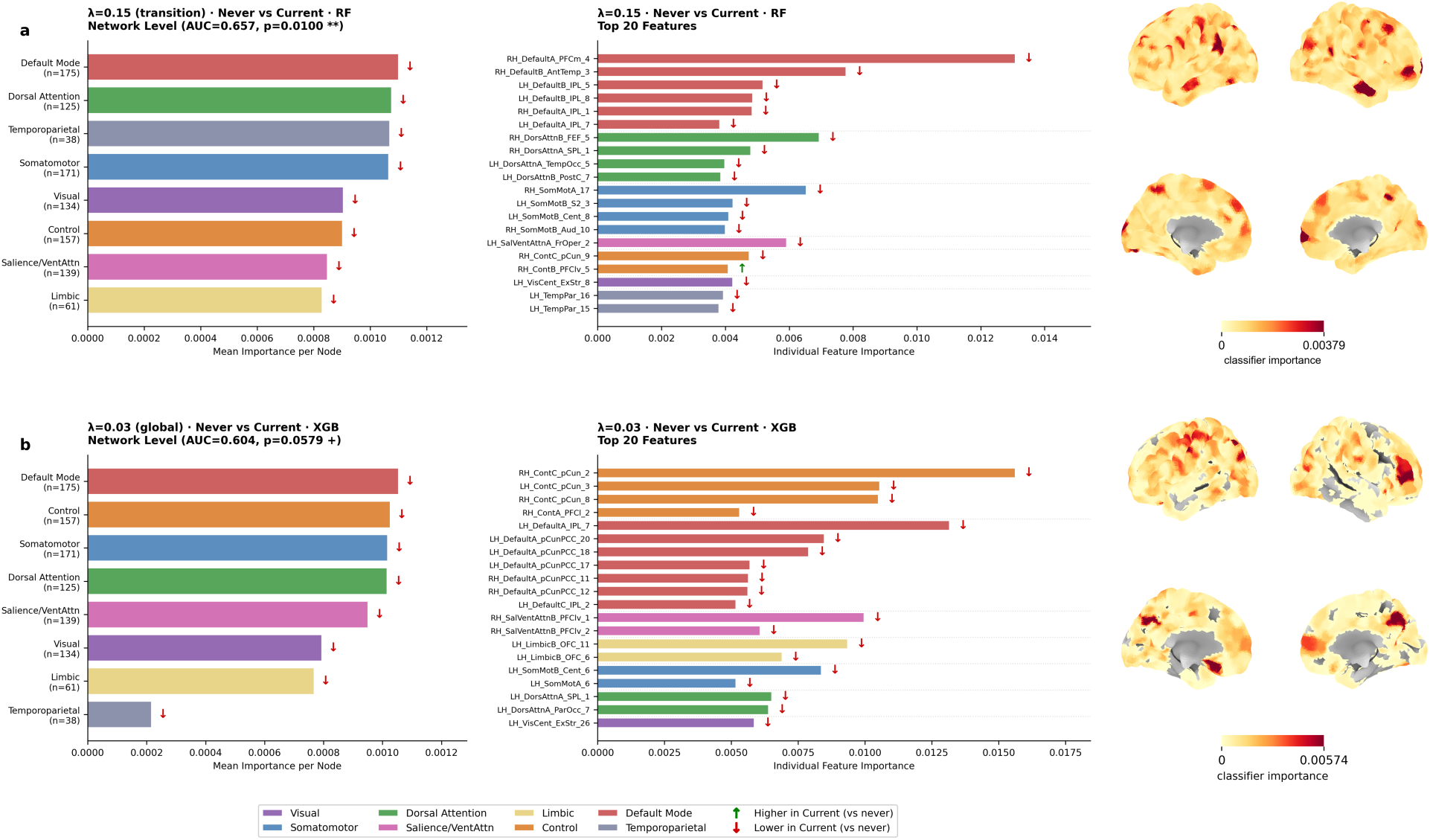
Network-level signature distinguishing never-users from current users. For each scale at which the never vs current comparison was significant or strong trend-level (*λ* = 0.15, *p* = 0.01; *λ* = 0.03, *p* = 0.0579, best model per scale) mean per-node feature importance aggregated within the eight cortical networks (left) and the twenty highest-ranked node features (right), with direction arrows (higher/lower in current users vs never-users). At *λ* = 0.15 the right medial-prefrontal DMN node (RH DefaultA PFCm) ranks first, with a profile of reduced nodal turbulence in users relative to never-users. Cortical surface renders of the per-node importance magnitude are shown alongside each row.

For long-term users (Figure 5), the discriminative signal was likewise distributed across all networks, with reduced nodal turbulence in users relative to never-users. The single most important feature was again the right medial prefrontal DMN node (RH DefaultA PFCm), which ranked first at all three significant scales (*λ* = 0.30, 0.18, 0.15), giving a consistent DMN medial-prefrontal anchor across the spatial hierarchy.

**Figure 5:**
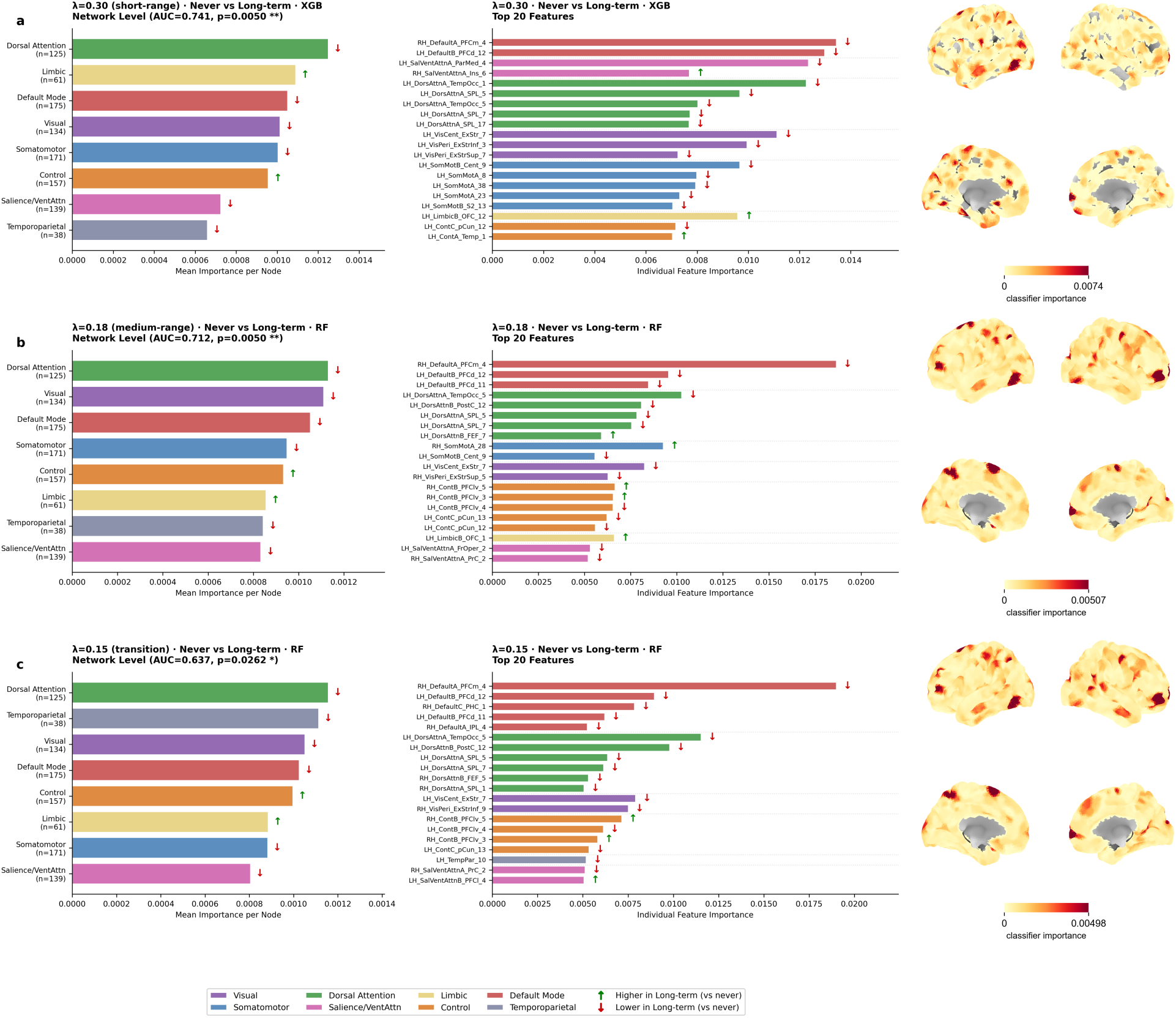
Network-level signature distinguishing never-users from long-term users. For each scale at which the never vs long-term comparison was significant (*λ* = 0.30, *p* = 0.005; *λ* = 0.18, *p* = 0.005; *λ* = 0.15, *p* = 0.0262; best significant model per scale), mean per-node feature importance aggregated within the eight cortical networks (normalized for network size; left) and the twenty highest-ranked individual node features (right), with arrows indicating the direction of the group difference (higher/lower in long-term users vs never-users). A right medial-prefrontal DMN node (RH DefaultA PFCm) ranks first across all three scales. Cortical surface renders of the per-node importance magnitude are shown alongside each row.

At the network level, the dorsal-attention system ranked highest across scales, followed by varying combinations of the DMN, visual, limbic, and temporoparietal systems. Importance-weighted nodal turbulence was reduced in long-term users across most leading networks, but the limbic, control, and salience networks moved in the opposite direction, showing increased nodal turbulence in long-term users relative to never-users, consistently across short and long distance scales (*λ* = 0.30, 0.18, 0.15). Among the twenty most important nodes at *λ* = 0.30, this increase was carried by a left orbitofrontal limbic node (LH LimbicB OFC), a left temporal control node (LH ContA Temp), and a right insular salience/ventral-attention node (RH SalVentAttnA Ins), all elevated in longterm users, while the leading DMN, dorsal-attention, and visual nodes were reduced. The leading node and several leading networks shared between the current-user and long-term signatures at *λ* = 0.15 suggest that the separability emerging in current users at the transition reflects the same distributed, DMN-anchored signature that characterizes cumulative exposure, detectable even without long duration.

### Brain dynamics reorganize following oral contraceptive cessation

Past users (*N* = 74) showed a change in the direction of association across most metrics relative to current users (Figure 6), suggesting active post-cessation remodeling rather than a passive return to baseline. Brain Turbulence showed a directional reversal that did not reach significance (*p >* 0.37). Information Transfer was positively associated with prior duration across 8 of 11 scales (*β* ≈ +0.31; e.g., *λ* = 0.30: *p* = 0.010), and Information Cascade Flow was negatively associated across 8 of 10 scales (*β* ≈ −0.28; e.g., *λ* = 0.15: *p* = 0.018). The global Information Cascade similarly reversed (*β* = −0.271, *p* = 0.028). The magnitude of reorganization varied with prior duration rather than converging on the never-user profile. While the absence of time-since-discontinuation data limits our ability to disentangle prior exposure from recovery time, this scale-spanning reversal suggests active remodeling of brain dynamics following cessation. The same dissociation appeared at the multivariate level: past users were separable from current users (RF AUC = 0.63, *p* = 0.028; XGB AUC = 0.63, *p* = 0.024) and from long-term users (RF AUC = 0.65, *p* = 0.034, at the short scale *λ* = 0.30), but remained near chance against never-users (AUC ≈ 0.59). Past users thus resemble never-users more than they resemble women currently taking the pill, consistent with a partial return toward the never-user profile after discontinuation, even as the sign-reversed, duration-scaled association marks the post-cessation state as actively remodeled rather than fully normalized.

**Figure 6:**
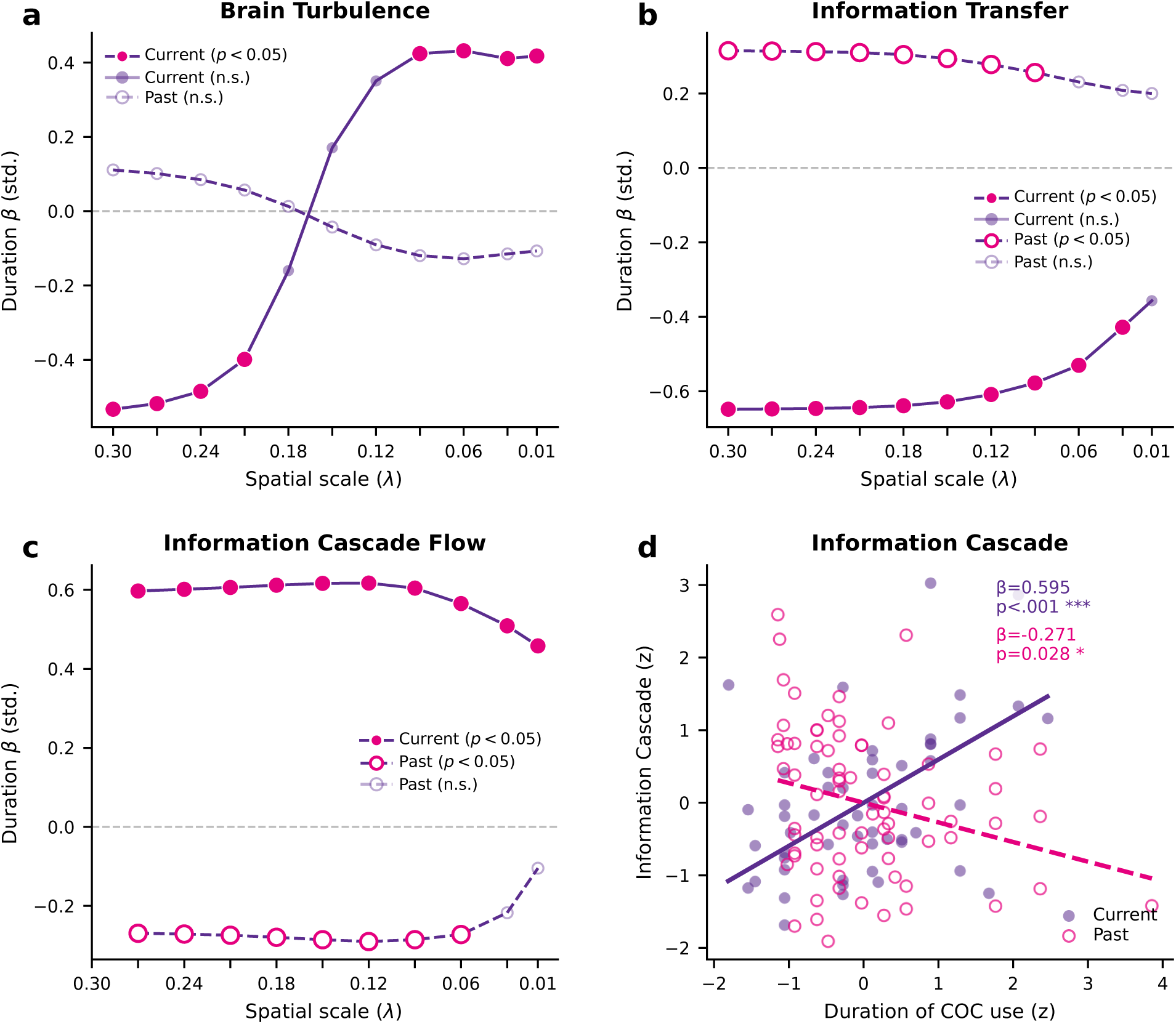
Brain dynamics reorganize following oral contraceptive cessation. Scale-level regression coefficients (*β*, standardized) for the association between prior COC duration and brain dynamics in current (*N* = 54, solid line) and past users (*N* = 74, dashed line), controlled for age. **(a)** Brain Turbulence shows directional reversal across scales without reaching significance in past users (*p >* 0.37). **(b)** Information Transfer reverses from negative (current) to positive (past; *β* ≈ +0.31; e.g., *λ* = 0.30: *p* = 0.010), significant at 8 of 11 scales. **(c)** Information Cascade Flow reverses from positive (current) to negative (past; *β* ≈ −0.28; e.g., *λ* = 0.15: *p* = 0.018). **(d)** Global Information Cascade reverses sign in past users (*β* = −0.271, *p* = 0.028) relative to current users (*β* = 0.595, *p <* 0.001). Filled circles: current users (*p <* 0.05); open circles: past users (*p <* 0.05).

## Discussion

Our findings suggest that cumulative oral contraceptive (COC) exposure is associated with duration-dependent, age-independent differences in whole-brain turbulent dynamics. This pattern is characterized by a shift from local to global communication, where longer exposure suppresses local synchronization and spatial information transfer while accelerating cross-scale hierarchical cascades. Machine learning pipelines supported the robustness of this duration signature, predicting individual years of use and accurately classifying active and long-term users from never-users across multiple spatial scales. The underlying predictive topography was widely distributed across multiple large-scale networks, led by the DMN in current users and by dorsal-attention in long-term users, with limbic and control systems reversing direction under long-term exposure. Finally, past users exhibited a scale-spanning sign reversal that scaled with historical duration rather than returning to a never-user baseline. Together, these results characterize cumulative COC use as a neuroendocrine modulator of multiscale brain communication.

Longer exposure was associated with a shift in the balance between local synchronization and structured cross-scale flow, characterized by lower local information transfer and higher hierarchical cascade flow. Within the turbulence framework this balance reflects how processing is organized across spatial scales (Deco and Kringelbach, 2020; Escrichs et al., 2022). Age did not predict any metric, despite being correlated with duration, suggesting an exposure-related rather than maturational effect in this age range. This cumulative shift parallels the acute hormonal modulation of turbulent dynamics described with the same framework across the natural menstrual cycle: in a dense-sampling study of a single woman scanned daily, the luteal phase showed higher long-distance turbulence and enhanced transfer and cascade flow relative to the follicular phase, and a hormonal-contraceptive regime abolished these cyclic shifts to yield a flattened, stable profile (De Filippi et al., 2021).

The ability to predict individual duration of use from brain dynamics beyond age suggests that this reorganization follows a consistent pattern rather than random variation. The predictive gain was carried by information transfer and cascade flow, measures of the spatial and cross-scale propagation of synchronization, rather than by any single focal metric, which is consistent with cumulative exposure reshaping distributed communication rather than an isolated region. A single-subject dense-sample study pointed in the same direction: oral contraceptive use lowered network modularity, system segregation, and characteristic path length relative to the natural cycle, indicating a less segregated, more globally connected architecture (Jensen et al., 2022).

The machine-learning approach suggests that this reorganization is systemic rather than localized. Individual cumulative exposure was decoded from brain dynamics across spatial scales. Because whole-brain turbulence captures the entire hierarchy of information processing (Deco and Kringelbach, 2020; Deco et al., 2025), this predictive capacity suggests that synthetic steroid exposure is associated with variation in the balance between local segregation and global integration (Beltz and Moser, 2020; Haase et al., 2025). Furthermore, the classification results show that never-users are highly discriminable from long-term users, across short and middle scales, where the cumulative signal was most pronounced. By contrast, including all current users was separable from never-users at the transitional scale (*λ* = 0.15) and showed a strong trend at the global scale (*λ* = 0.03), indicating that the observed dynamic reconfigurations reflect cumulative hormonal exposure. This dose-dependent pattern is consistent with previous works showing that resting-state effects of oral contraceptives are network-specific and relate to use history rather than forming a simple user/non-user dichotomy (Pletzer et al., 2016; Sharma et al., 2020).

Feature importance analysis revealed that the discriminative signal was distributed across cortical networks yet consistently led by a right medial-prefrontal DMN node, with dorsal-attention and DMN systems ranking highest at the network level in both long-term and current users. While most leading networks showed reduced nodal turbulence in users relative to never-users, the limbic and control networks moved in the opposite direction, with increased nodal turbulence in longterm users, carried by orbitofrontal (OFC) limbic, temporal control, and insular salience nodes. The orbitofrontal cortex is known to be highly sensitive to oral contraceptives in both structural and functional studies (Petersen et al., 2015; Gingnell et al., 2016). The involvement of the DMN and dorsal-attention systems is consistent with the node-level localization of cycle-dependent turbulence, where the DMN showed the largest modulation (De Filippi et al., 2021). It also aligns with the broader resting-state literature implicating the DMN and prefrontal cortex in hormonal-contraceptive and menstrual-cycle effects, including reduced DMN and anterior-cingulate connectivity in current users (Pletzer et al., 2016; Hidalgo-Lopez et al., 2023b,a).

In past users, the duration associations for information transfer and cascade flow reversed in sign relative to current users and scaled with prior duration, rather than returning to the never-user profile. Node-level classifiers refined this: past users were separable from current users and from long-term users, but not from never-users, indicating that the post-cessation state has moved closer to the never-user profile than to that of women currently taking the pill. The two observations are complementary: a graded, duration-scaled footprint of prior exposure persists and is recoverable by regression and by classification against current users, while the resemblance to never-users suggests a partial return toward the unexposed profile after discontinuation. A persisting, duration-related effect that nonetheless trends back toward baseline is consistent with evidence that some contraceptive-related effects outlast use: structural changes that remodel after cessation (Pletzer et al., 2019), and duration-dependent effects on verbal performance and associated activation that persist in past users (Noachtar et al., 2022).

Some limitations warrant consideration. The cross-sectional design cannot establish causality. Longitudinal studies following users from oral contraceptive initiation are required to characterize the trajectory of these reconfigurations. Second, the relatively narrow age range of our cohort highlights the need for future research that includes older populations with longer cumulative oral contraceptive use. Finally, the absence of time-since-discontinuation data in past users limits our ability to fully disentangle prior exposure duration from recovery time, and should be addressed in future work.

Taken together, these results suggest that prolonged oral contraceptive use induces a neuroendocrine reconfiguration of brain dynamics. The convergence of independent lines of evidence, the scale-dependent shift from local to global synchronization, the prediction of individual exposure duration from brain dynamics across scales, and the reversal of these signatures in past users suggest that cumulative hormonal exposure reshapes brain dynamics in a continuous, duration-dependent way. These results open a path toward individualized markers of brain health and provide a quantitative basis for tracking how brain dynamics evolve across the reproductive lifespan.

## Methods

### Participants

The sample comprised 192 women from a single acquisition site (TR = 2.25 s): 54 current COC users (age: 22.1 ± 2.7 years; duration: 4.7 ± 2.6 years, range 0.08–11.0), 78 past users (age: 25.1 ± 4.3 years; prior duration: 4.1 ± 3.4 years, available for 74 of 78), and 60 never-users (age: 22.7 ± 3.4 years; all scanned in the follicular phase). Data were collected across sub-studies with identical acquisition parameters and harmonized using CovBat (Chen et al., 2022). Of the current users, 52 had documented progestin formulation (androgenic, *n* = 31; anti-androgenic, *n* = 20). Participants had no history of psychiatric, neurological, or endocrine disorders and were free from concurrent medications. All procedures were approved by the local ethics committee of the University of Salzburg and conformed to the Declaration of Helsinki. All participants provided written informed consent.

### MRI acquisition and preprocessing

Resting-state functional MRI data were acquired on a Siemens TIM Trio scanner following a field map acquisition. Resting-state BOLD images were obtained using a T2^∗^-weighted gradient echo-planar imaging (EPI) sequence with 36 transversal slices oriented parallel to the AC-PC line, ensuring whole-brain coverage (TR = 2,250 ms, 216 volumes, TE = 30 ms, flip angle = 70^◦^, slice thickness = 3.0 mm, matrix = 192 × 192, FOV = 192 mm). To allow for magnetic field equilibration, the initial six volumes of each functional run were discarded. The remaining time series were despiked to mitigate extreme signal transients using AFNI’s 3dDespike utility. Subsequent spatial preprocessing was performed in SPM12. Functional volumes were realigned and unwarped utilizing field map data, and subsequently coregistered to their corresponding high-resolution structural images. Anatomical segmentation and spatial normalization to MNI space were performed using the CAT12 toolbox, with the resulting deformation fields applied to the functional data before spatial smoothing with a 6-mm full-width-at-half-maximum (FWHM) Gaussian kernel. Susceptibility-induced distortions were corrected by estimating field maps from paired EPI acquisitions with reversed phase-encoding directions using FSL’s topup tool (Andersson et al., 2003), from which voxel displacement maps were generated via the SPM12 FieldMap toolbox (Chang and Fitzpatrick, 1992). Rigorous quality control included the strict exclusion of participants with excessive head motion (exceeding 3 mm in translation or 2^◦^ in rotation) and visual verification of coregistration and normalization accuracy against standard T1 and EPI templates. To further isolate the underlying neural signal, motion-induced spurious variance was filtered out using the non-aggressive mode of independent component analysis-based automatic removal of motion artifacts (ICA-AROMA) (Pruim et al., 2015). Finally, mean regional BOLD time series were extracted for each participant utilizing the 1,000-node Schaefer cortical parcellation (Schaefer et al., 2018). BOLD time series were bandpass-filtered (0.008–0.08 Hz, second-order Butterworth filter).

### Data Harmonization

To account for variability between the sub-studies, biological features were harmonized using Cov-Bat (Chen et al., 2022). CovBat adjusts both the mean and the covariance of imaging features across acquisition sites. To ensure that the biological variance of interest was preserved during this process, biological age, duration of COC use, and subgroup were included as protected covariates in the harmonization model.

### Brain dynamics mathematical formulation

The mathematical formalization of brain turbulence, including the computation of local Kuramoto order parameters, spatial information transfer, and cross-scale information cascade flow, was carried out as in (Escrichs et al., 2022). The local Kuramoto order parameter was calculated as:

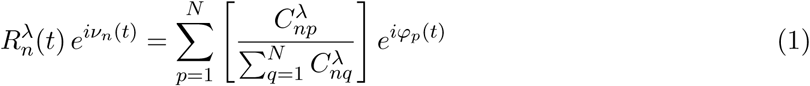

where *φ_p_*(*t*) is the instantaneous BOLD phase of region *p* (Hilbert transform of filtered signal), and *C^λ^* is a distance-dependent coupling kernel controlled by *λ*. High *λ* (= 0.30) restricts coupling to nearby regions; low *λ* (= 0.01) allows coupling across large distances. Eleven scales were computed: *λ* ∈ {0.30, 0.27, . . ., 0.01}.

Turbulence was defined as:

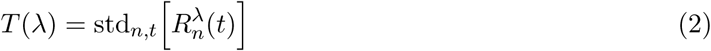

Spatial information transfer was calculated as the log-log slope of the temporal correlation between region pairs as a function of Euclidean distance:

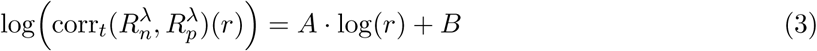

Information cascade flow was calculated as:

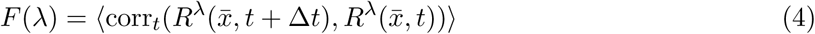

Finally, the information cascade was defined as the mean of *F* (*λ*) across all scales.

### Duration regression analyses

The association between cumulative COC use and each brain-dynamics metric was assessed with least-squares regression, modelling each metric at each spatial scale as Metric(*λ*) ∼ Duration + Age with all variables z-standardized, so that coefficients are expressed in standard-deviation units, the global Information Cascade, a single scalar, was modelled analogously. Models were fitted separately for current and past users. Because chronological age and duration of use were collinear (current users: *r* = 0.676, VIF= 1.84; past users: *r* = 0.320, VIF= 1.11; both well below the threshold of concern, VIF*<* 5), the variance inflation factor was computed and reported so that age-adjusted coefficients can be interpreted accordingly. To test whether the duration slope differed between current and past users, a pooled interaction model Metric ∼ Duration × Group + Age (Group: current vs. past) was fitted on pooled z-standardized predictors, with the Duration×Group term quantifying the slope difference; a sign reversal was considered supported only when this interaction was significant and both within-group slopes were individually significant with opposite sign, otherwise a mere change of sign is reported descriptively.

### Individual duration prediction

To assess whether brain metrics could predict individual oral contraceptive (COC) duration, we implemented a machine-learning pipeline based on Lasso regression (Tibshirani, 1996). Given the inherent collinearity between chronological age and years of use, we isolated the unique predictive power of the brain metrics beyond the effects of aging by comparing nested models that include age as a covariate (the Δ*r* approach described below), rather than by residualizing age out of the features. Model performance was evaluated using a nested 10-fold cross-validation (CV) scheme to prevent data leakage. Within each outer training fold, features were standardized, and a LassoCV model with a 3-fold inner validation was employed to optimize the regularization hyperparameter (*α*). The trained model was subsequently applied to predict the duration of use in the unseen test fold. For each brain metric and across all spatial scales (*λ*), two competing models were evaluated under the identical nested CV scheme: a *base model* using only chronological age, yielding a baseline out-of-fold Pearson correlation (*r*_base_), and a *full model* combining age and the corresponding brain metric (*r*_full_). The unique predictive capacity of the brain dynamics was quantified as Δ*r* = *r*_full_ − *r*_base_. Statistical significance was assessed via conditional permutation testing (1000 permutations), in which metric values were randomly shuffled across subjects while keeping the age-duration relationship intact. The empirical *p*-value was computed with the (*k* + 1)*/*(*n* + 1) estimator (Phipson et al., 2010), with significance set at *p <* 0.05. No correction across scales was applied, as they are points of a single continuous spatial decomposition of the same metric rather than independent tests.

### Multiscale group classification and reference-scale inference

To evaluate whether multi-feature patterns of brain dynamics could distinguish binary groups, we used an age-residualized multivariate classification pipeline on a high-dimensional feature space (1033 features: 1000 regional node-level turbulence values plus 33 global descriptors spanning turbulence, information transfer, and cascade flow), separating never-users from COC users (never vs current; never vs long-term, duration ≥ cohort median, 4.75 y). The pipeline had two sequential stages. First, an exploratory *scale sweep* was run across all 11 spatial scales (*λ* = 0.30 to 0.01) using a single-permutation framework to map the descriptive ROC AUC landscape without formal inference. Second, guided by this landscape, inference was performed at the scales where the discriminative signal was concentrated. Because the spatial scales are points of a single continuous decomposition of the same metric rather than independent tests, this targeted inference reads where on the continuum the signal peaks rather than selecting among competing hypotheses; concentrating the permutation testing on the relevant scale regime also avoids the substantial computational cost of full permutation across every scale. For the principal never-versus-OC analyses, four reference scales spanning the hierarchy were used (short, *λ* = 0.30; middle, *λ* = 0.18, 0.15; global, *λ* = 0.03). For the past-user comparisons, where the descriptive sweep identified the peak discriminative scale for each classifier, inference was directed to those peak scales. At each reference scale, classification used two complementary models: Random Forest (RF) (Breiman, 2001) and Extreme Gradient Boosting (XGBoost) (Chen and Guestrin, 2016), embedded in an outer 10-fold stratified cross-validation loop. Within each outer fold, a Ridge regression (Hoerl and Kennard, 1970) predicting each feature from chronological age was fitted only to the training partition and then applied to remove the linear age component from both the training and test sets, neutralizing age-related leakage. Class imbalance was addressed with the Synthetic Minority Over-sampling Technique (SMOTE) (Chawla et al., 2002) strictly within the training fold. Hyperparameters for both classifiers were optimized via an inner 3-fold randomized search (*n*_iter_ = 10) to maximize ROC AUC. Significance at the reference scales was determined by label permutation tests (*N*_max_ = 1000) with a symmetric adaptive early-stopping rule defined *a priori* : a null stop was triggered after *m* ≥ 100 permutations if the lower 95% bound of the Monte-Carlo *p* exceeded 0.30, and a success stop after *m* ≥ 200 permutations if the upper 99% bound fell below *α* = 0.05. Exact empirical *p*-values were computed with the (*k*_exc_ + 1)*/*(*m* + 1) estimator (Phipson et al., 2010), preventing zero-valued probabilities and ensuring consistency with the early-stopping boundaries. Performance is reported as age-residualized ROC AUC.

### Feature-importance profile and cortical mapping

To characterize the substrates driving classification, feature-importance profiles were extracted from the best-performing significant model at each reference scale, using impurity-based importance for Random Forest and gain-based importance for XGBoost. To obtain stable estimates, importances were averaged across the ten cross-validation folds, each fitted under the same within-fold age-residualization and SMOTE pipeline used for classification. Because the feature space contained both nodal and global variables, spatial localization was performed on the 1000 cortical nodes (Schaefer et al., 2018). To identify network-level patterns, the unsigned per-node importance magnitudes were averaged within the eight canonical functional networks (normalized for network size). To capture regional specificity, the 20 highest-ranked individual node features were isolated, and the sign of the group-level difference (mean in COC users minus never-users) determined the direction of each effect. Finally, the 1000-node importance vector was projected onto a cortical surface to visualize the spatial topography of the predictive signature as an unsigned magnitude map.

## Acknowledgements

A.E. was supported by the European Union’s Horizon Europe research and innovation programme under the Marie Sklodowska-Curie Actions (ID: 101207460, NEUROCONTRA, HORIZON-MSCA 2024-PF-01-01), by the José Castillejo Mobility Grant (CAS23/00099), Spanish Ministry of Science and Innovation, and by the project eBRAIN-Health (ID: 101058516) funded by EU Horizon Europe.

Y.S.P. and G.D. were supported by the project NEMESIS (ref. 101071900) funded by the EU ERC Synergy Horizon Europe. Y.S.P. was also supported by the Grant PID2024-162576NA-I00 funded by MICIU/AEI/10.13039/501100011033 and by “ERDF A way of making Europe”. B.P. was supported by the European Research Council (ERC) Starting Grant 850953. A.E. and B.P. were also supported by the project NEUROFEM, funded by La Fundacío la Marató de TV3 (ID: 202410-30-31).

## Author contributions

A.E.: conceptualization, methodology, data analysis, writing original draft. Y.S.P: methodology, review & editing. G.D.: methodology, review & editing. B.P.: conceptualization, data collection, review & editing.

## Code availability

The code supporting this work will be available on GitHub upon acceptance: https://github.com/aescrichs/pill.

